# Cryo-electron microscopy of cytoskeletal ‘fibril’ involved in *Spiroplasma* swimming

**DOI:** 10.1101/2023.06.28.546849

**Authors:** Yuya Sasajima, Takayuki Kato, Tomoko Miyata, Hana Kiyama, Akihiro Kawamoto, Fumiaki Makino, Keiichi Namba, Makoto Miyata

## Abstract

*Spiroplasma*, parasitic or commensal bacteria, can swim by switching the handedness of its helical cell body. A helical cell body is formed by an internal ribbon of MreB, an actin superfamily, and *Spiroplasma*-specific fibril proteins. Here we have solved the structure of the fibril filament by single-particle cryo-electron microscopy at 3.6 Å resolution and built its atomic structure. The structure is composed of repeated rings and cylinders. The N-terminal cylinder of the fibril protein shows a structure similar to that of S-adenosylhomocysteine nucleosidase, while the C-terminal ring does not show similarity to other proteins. The filament is nonpolar and flexible, possessing a helical pitch of 700 nm, consistent with cell helicity. Cryo-electron tomography revealed aligned several MreB filaments in the center of the ribbon, flanked by membrane-binding fibril filaments through electrostatic interactions. This study discusses the evolution and roles of the fibril filament.

## Introduction

*Spiroplasma* sp., a member of the class Mollicutes characterized by a small genome size, is spread worldwide (*1*). They significantly impact industry and organisms other than themselves, including plants such as corn and citrus fruits and arthropods such as crabs and flies. One of the successful features of this bacterium may be its unique swimming mechanism, leading to efficient migration and, thus, habitat expansion (*2*). Swimming is facilitated by switching the helical cell body between left- and right-handed, in which the rotation of the cell helix behind a kink caused by the switching of helix handedness pushes the water backward (*3-6*). The helix formation and switching in *Spiroplasma* cells lacking a peptidoglycan layer are plausibly caused by an internal “ribbon” structure (*7, 8*). The ribbon is composed of MreB (*8-10*), known as bacterial actin (*11*), and fibril, a protein specific to the genus *Spiroplasma* (*12, 13*). A recent study reported that combining two types of *Spiroplasma* MreB proteins causes motility similar to swimming in a minimal synthetic bacterium, JCVI-syn3B (syn3B) (*14-16*). This indicates that the conformational change in MreB is responsible for the force generation for swimming. MreB is a bacterial actin that has a critical role in peptidoglycan synthesis in walled bacteria (*17*) and diverges into five classes in the evolution to class Mollicutes (*18, 19*). The fibril protein, another significant component of the ribbon, is one of the most abundant proteins in *Spiroplasma* cells and is well conserved in the genus *Spiroplasma*, as featured by more than 90% amino acid sequence similarity (*3, 9, 12*). Interestingly, the helical pitch of the isolated fibril filaments is comparable to that of a *Spiroplasma* cell. Moreover, the N-terminal half of the fibril protein shares up to 20% similarity with S-adenosylhomocysteine (SAH) nucleosidase, which is essential for bacterial growth (*3, 7, 20*). These points raise questions about the possible role and evolutionary origin of the fibril protein in *Spiroplasma* swimming and survival. To answer these questions, determining the structure of the fibril protein and the ribbon construction is crucial. Previously, we isolated the fibril protein from *Spiroplasma* cells and showed a low-resolution structure (*12*). In this study, we performed single-particle cryo-electron microscopy (cryo-EM) of the isolated fibril filament and cryo-electron tomography (cryo-ET) of *Spiroplasma* and the motile syn3B cells to clarify the fibril protein.

## Results

### Cryo-EM single-particle analysis of the fibril filament

We isolated fibril filaments from *Spiroplasma* cells using a previously established method (*12*) and performed cryo-EM observation for single particle analysis. As good-quality images were obtained (Fig. 1a), segmented images of 300 nm length along the filaments were extracted with 90% overlap between consecutive segments, followed by 2D classification (Fig. 1b). The 3D image was then reconstructed for the single-stranded filament from 13,825 segmented images. The resulting structure had a resolution of 3.6 Å with a Fourier shell correlation of 0.143 (Supplementary Fig. 1 and 2). Based on the cryo-EM map, including six subunits, an atomic model of the fibril protein was constructed (Fig. 1c-d). To estimate the helical pitch and handedness of the fibril filament, we made a long 3D density map of the fibril filaments by extending the short segment map to a length of half helical pitch (Fig. 1e). The reconstructed fibril filament was left-handed, and the half-helical pitch was approximately 350 nm. The shape of the fibril filament was characterized by the rings with a 105 Å long axis, connected by the cylinders with an 89 Å periodicity (Fig. 1c). These characteristics of the fibril filament were consistent with the structure previously revealed by negative staining electron microscopy (*12*).

**Fig. 1.**
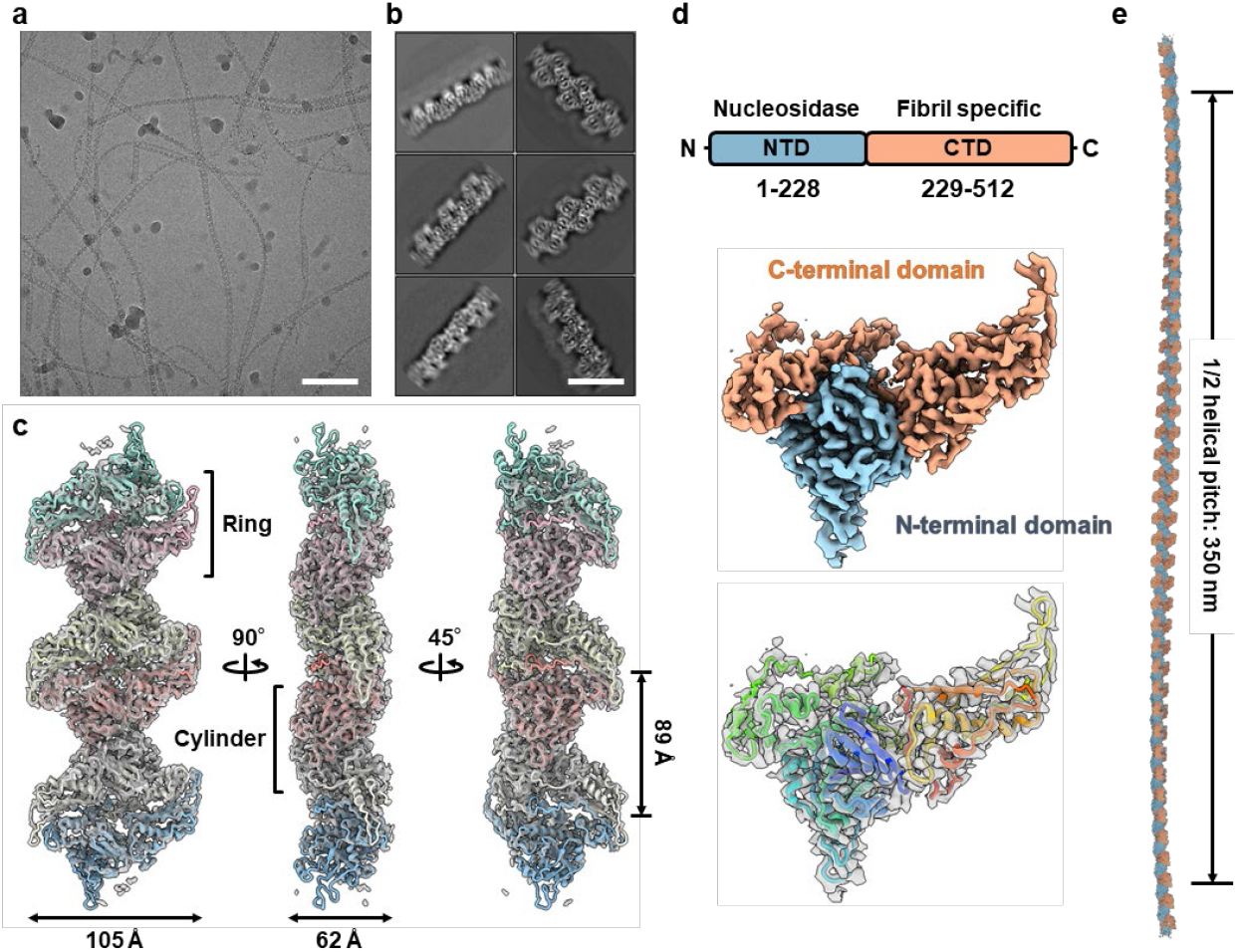
Single-particle cryo-EM of the fibril filament. **a** A cryo-EM image of ice-embedded fibril filaments. Scale bar, 100 nm. **b** Class averages by 2D classification. Scale bar, 15 nm. **c** Reconstructed density map and atomic models of the fibril filament. Each fibril subunit is colored differently. **d** (Upper) Domain schematic of the fibril protein. (Middle) EM density map of a fibril subunit. The N-terminal domain (NTD) and C-terminal domain (CTD) are colored blue and orange, respectively. (Lower) An atomic model in ribbon format colored rainbow from the N-to C-termini is overlaid to the EM density map. **e** A density map of the fibril filament extended to the length of half helical pitch.

### Atomic model of the fibril filament

In the atomic model, all 512 amino acid residues of the fibril protein were assigned to the cryo-EM map (Fig. 1d, Supplementary Fig. 2). The amino acid sequence of the fibril protein can be divided into the N-terminal 228-residue sequence with approximately 20% similarity to SAH nucleosidase and the C-terminal sequence. The atomic model of the fibril filaments showed that the 228 residues on the N-terminal side mainly correspond to the cylinder, whereas the C-terminal region primarily corresponds to the ring (Fig. 1d).

Three interaction surfaces (i) -(iii) were found between subunits in the fibril filament, which were characterized by the dominance of non-polar side chains (Fig. 2a). The interface (i) consists of three α-helices, including four non-polar and three polar side chains. Interface (ii) consists of four loops, including eight non-polar and two polar side chains. Interface (iii) consists of four β-sheets and one loop, including 13 non-polar side chains. These interactions may suggest elasticity and robustness as characteristics of fibril filaments.

**Fig. 2.**
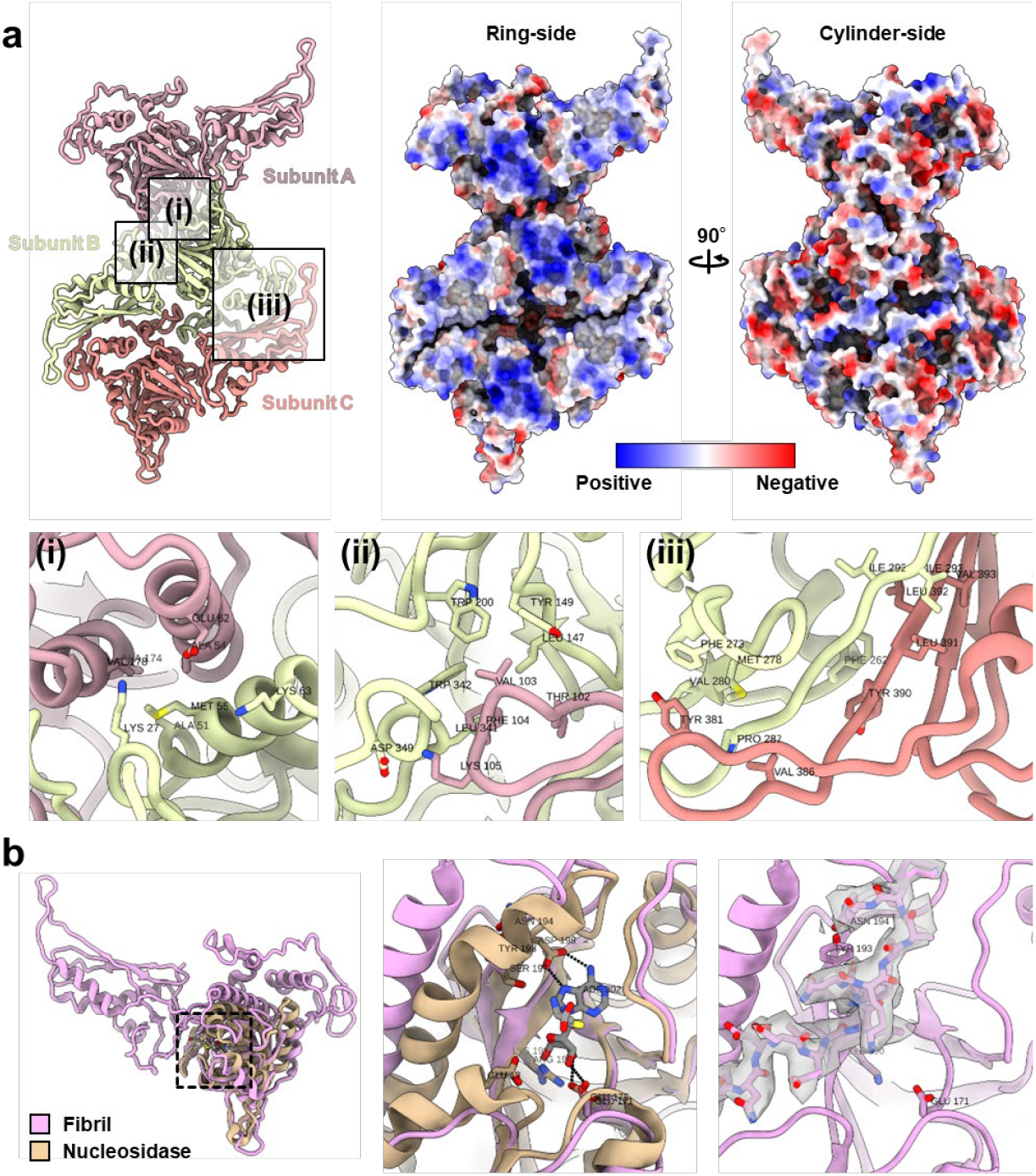
Atomic structure of fibril filament. **a** (Upper left) Atomic structures of the fibril filament composed of three subunits. The areas enclosed by squares (i)–(iii) are magnified below. Note that the magnified images are viewed from different angles. (Upper right) Surface model colored for electrostatic potential at pH 7.0. **b** Superposing of the fibril protein (pink) and the nucleosidase (beige). The boxed area is magnified in the right panels. The right panel shows a part of the EM density map as translucent density.

We compared the fibril protein structure with that of SAH nucleosidase from *Escherichia coli* (IJYS) (Fig. 2b, Supplementary Fig. 3) (*21*). In SAH nucleosidase, the adenine-binding site consists of Glu175, Arg194, Ser197, and Asp198 on two β-strands, an α-helix, and a loop. In contrast, the residues of the fibril protein corresponding to the adenine binding site of the nucleosidase were partially mutated to Glu171, Lys190, Tyr193, and Asn197, respectively (Supplementary Fig. 3). In addition, the Val335–Lys347 chain of the C-terminal domain of the same subunit was found to bind to the position corresponding to the adenine-binding pocket (Fig. 2b, Supplementary Fig. 3). The α-helix composed of residues Met10–Gly22 of the nucleosidase is replaced by β-strands composed of residues Tyr9–Lys13 and Val333–Asn336 in the fibril protein (Supplementary Fig. 3). Based on these observations, we concluded that the fibril protein does not retain nucleosidase activity.

### Structural flexibility of the fibril filament

To discuss possible conformational changes of the fibril filament, we examined its structural variation in the cryo-EM particle images. The 147,040 particle images used to reconstruct the high-resolution cryo-EM map were re-extracted with a box size of 450 nm to include 10 fibril protein subunits. The 2D classification was performed based on these images to identify possible single-stranded filaments. The resulting 81,139 particles were subjected to 3D variability analysis in the cryoSPARC software. The structural changes of the fibril filaments were specified by continuous movements in three directions, while distinct movements were found in two directions. One was a forward bending toward the ring, and the other was a bending lateral to the ring (Fig. 3a–b). These flexibilities allow variations in the helical pitch and a diameter of 300–680 and 20–100 nm, respectively (Fig. 3c).

**Fig. 3.**
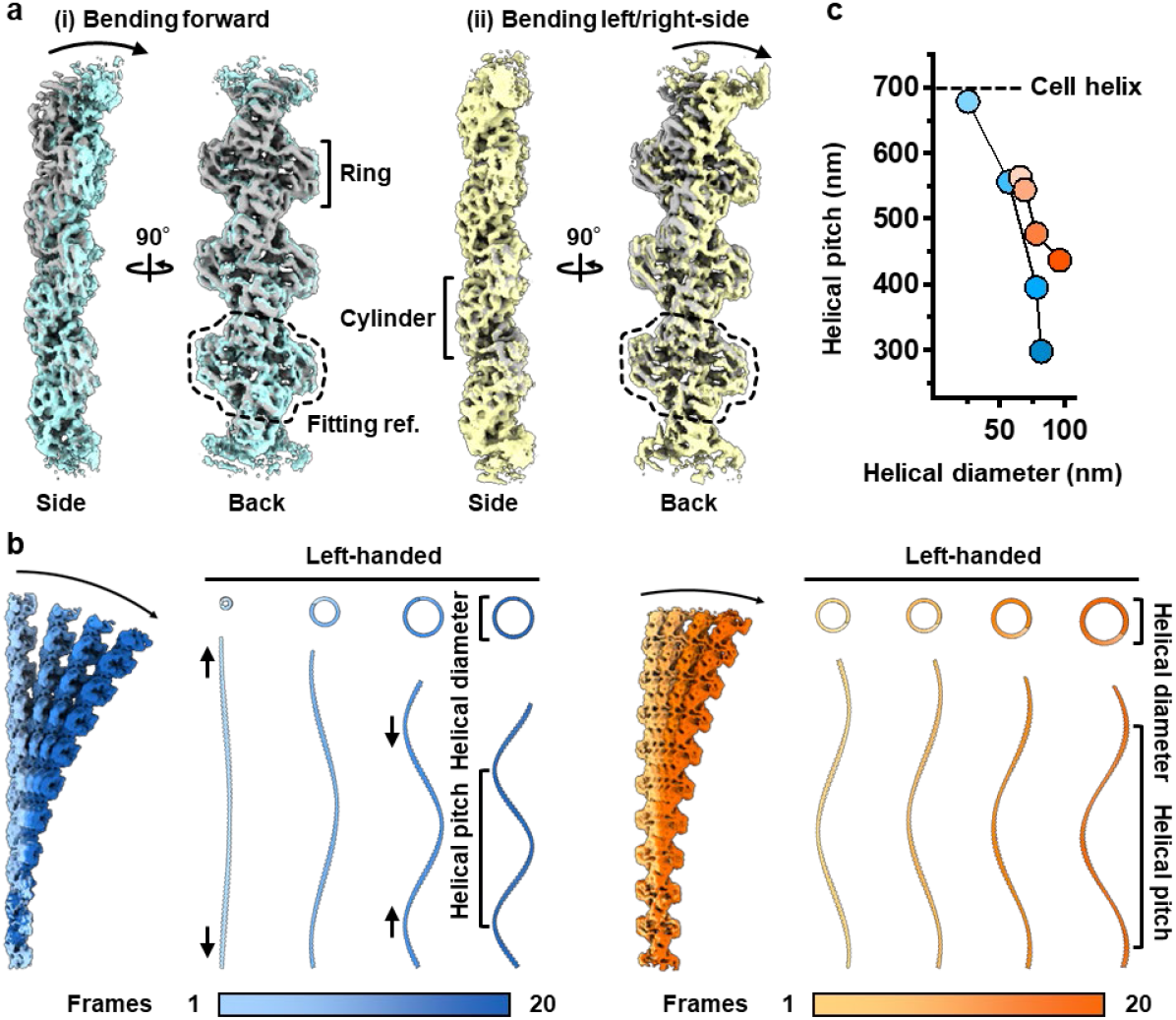
Structural flexibility of the fibril filament. **a** EM density map reconstructed by cryoSPARC 3d variability analysis. A default fibril filament (gray), bending to the ring side (blue), and bending to a lateral side (orange) are shown. The dashed area was used as a fitting reference. **b** Stitched volumes of straight and bent maps from a. Arrows indicate a spring-like extension and contraction of the fibril filament. Helical pitches and diameters corresponding to each frame are listed. **c** Relation between helical pitch and diameter.

### Cryo-ET of the internal ribbon

We examined the internal ribbon structure composed of MreB and fibril proteins by cryo-ET in *Spiroplasma* cells. *Spiroplasma* cells suspended in PBS were applied to EM grids, blotted from one side of the grid, and rapidly vitrified in liquid ethane. The prepared grids were imaged by cryo-EM in continuous tilt. The motion-corrected tilt series were aligned, and the three-dimensional tomograms were reconstructed. The reconstructed tomograms of the *Spiroplasma* cells visualized the internal ribbon structure running the whole length of the helical cell body in the more than 30 cells examined (Fig. 4a) (Movie S1). The ribbon was characterized as a sheet structure with several filaments in top view. It was bound to the cell membrane at its lateral edge in a sectional view at a middle height, suggesting that the ribbon is localized in the innermost line of the helical cell body (Fig. 4a). The cells showed both left and right helix handedness as well as a half-helical pitch distributing from 320 to 500 nm, which are consistent with those observed by optical microscopy, suggesting that the cells retained structural characters in the vitrified ice (Fig. 4b) (*12*). The inter-filament distance in the ribbon differed with the filament positions, i.e., approximately 10 and 5 nm in the central and peripheral parts, respectively (Fig. 4c).

**Fig. 4.**
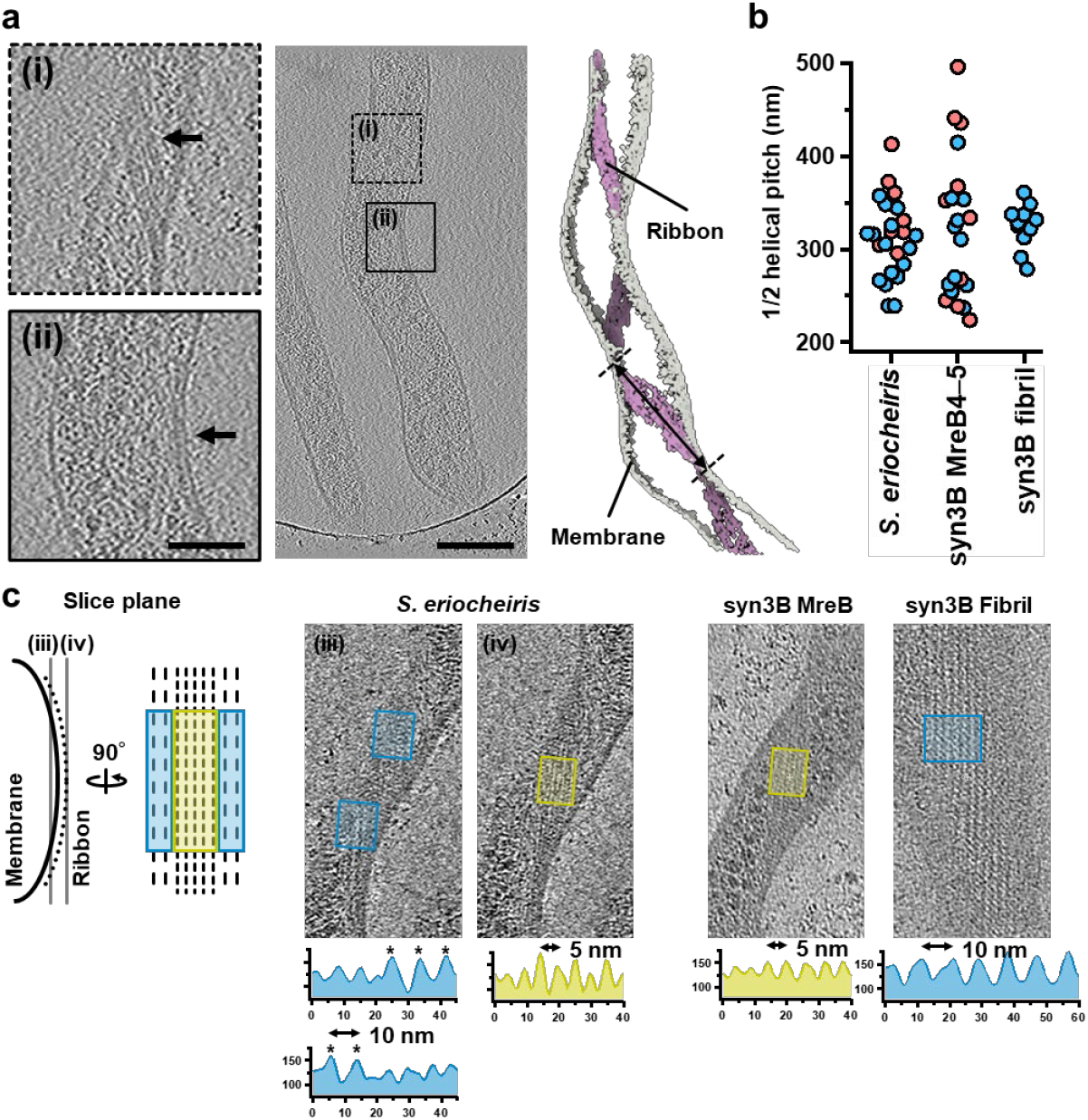
Cryo-ET of the internal ribbon structure. **a** Cryo-ET image of *Spiroplasma* cell. A slice image is shown in the center panel, and slice images of boxed areas (i) and (ii) are shown at different heights in the left panels. An arrow marks ribbons. Tomogram rendering is shown in the right panel. The bidirectional arrow indicates the half-helical pitch of the ribbon. **b** Distribution of half helical pitches of different cells. The results of right- and left-handed helices are colored red and blue, respectively. **c** Positions of two types of filamentous structures in internal ribbons. (Left) Schematic illustration for positions of two filamentous structures in a *Spiroplasma* cell. (Middle) Filamentous structures visualized in tomogram of a *Spiroplasma* cell. Density profiles are shown at the bottom. (Right) Filamentous structures visualized in tomogram of syn3B cells with MreB and fibril proteins. Density profiles are shown at the bottom. 10 or 5 nm peak intervals can characterize filamentous structures.

To identify the two filaments in the ribbon of *Spiroplasma* cells composed of MreB and fibril proteins, the syn3B cells expressing either a combination of MreB4 and MreB5 (MreB4–5) (*14*) or the fibril protein were subjected to cryo-ET. The ribbon-like sheet structures bound to the cell membrane were found in the syn3B cells of both constructs, running the whole length of the helical parts of the cell body in all of the more than 30 cells examined (Fig. 4c) (Movie S2). The half-helical pitch was approximately 300 nm (Fig. 4b). The syn3B cells with MreB4–5 showed both left and right helical handedness, while the syn3B cells with fibril protein showed only left helical handedness. These observations are consistent with previous optical microscopy observations (*12*). The peak intervals estimated from the protofilament density profiles were 5 and 10 nm for the cells expressing MreBs and fibril, respectively (Fig. 4c). Based on these observations, we concluded that the ribbon structure of *Spiroplasma* cells is composed of the MreB and fibril filaments in the central and peripheral parts, respectively (Fig. 4c).

### Subtomogram averaging of the *in situ* fibril filament

To clarify the intracellular structure of the fibril filament, we performed subtomogram averaging for cryo-ET of the syn3B cells expressing the fibril protein (Supplementary Fig. 4 and 5). The fibril filaments in the syn3B cells formed sheet-like structures in contact with the cell membrane (Fig. 5a). The density maps of the subtomograms containing three fibril filaments were manually extracted from the tomograms. The extracted particles were then sorted by 2D and 3D classification, and finally, a density map was obtained at a resolution of 16 Å from 5182 particles. The obtained structure had rings facing the cell membrane and cylinders facing the cytoplasm, forming a sheet with adjacent fibril filaments (Fig. 5b–c). When docked to the density map, the atomic model of the fibril protein showed that the side chains at the interface of neighboring fibril filaments were positioned to partially neutralize their positive and negative charges in the C-terminal domain. (Fig. 5c). The interaction between the fibril filament and the cell membrane, specifically the ring portion, was characterized by 14 positively charged residues of the fibril protein. This suggests their electrostatic interactions with the phosphate head groups of the lipids. (*22-24*).

**Fig. 5:**
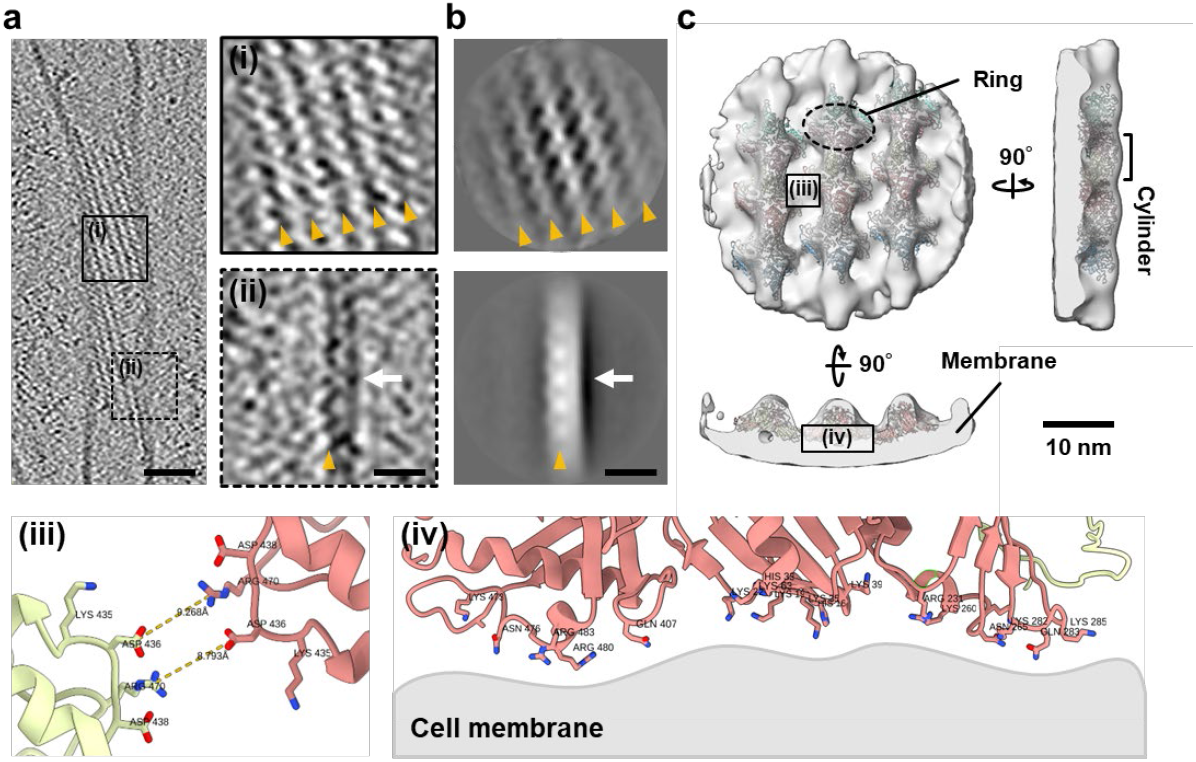
Subtomogram averaging of the *in situ* fibril filament. **a** Tomogram slice of syn3B cell expressing the fibril protein. The boxed square areas are magnified in the right panels. Solid and dotted squares show slices at different heights. Yellow triangles indicate individual fibril filaments. The white arrow indicates the cell membrane. **b** Projection images of subtomogram averages. Yellow triangles indicate individual fibril filaments. The white arrow indicates the cell membrane. **c** Subtomogram average of the *in-situ* fibril filament at 16 Å resolution. The atomic structures of the fibril filament fit the EM map. Individual fibril subunits are colored differently. The boxed areas (iii) and (iv) are enlarged at the bottom. Individual subunits are colored rose and yellow.

## Discussion

The N-terminal domain structure of the fibril protein supports the sequence analysis, suggesting that the fibril protein originated from a nucleosidase and achieved a completely different function (Fig. 2b) (*3, 7, 12*). The phylogenetic tree composed of this domain and the nucleosidase (Supplementary Fig. 3a), based on their amino acid sequences, unexpectedly suggested that the fibril protein is more closely related to other genera in the Mollicutes class than to the *Spiroplasma* genus. This may indicate that horizontal gene transfer was involved in fibril evolution (*25*). Regarding the C-terminal domain, we could not suggest an evolutionary scenario from the structure (Fig. 1d).

The structural flexibility was suggested in two directions (Fig. 3a, b). However, these changes are expected to be mixed in actual movements. Because the fibril filament was not depolymerized even by strong sonication (*12*), the fibril filament should have a flexible yet robust spring-like property that supports the helical shape of the cell. The proposed structural changes of the fibril filaments are continuous and do not involve distinct helicity switching (Fig. 3b). The helicity switching that propagates along the entire cell length during *Spiroplasma* swimming suggests a reversal of handedness in the fibril filaments (*3, 4, 6, 26*). However, the structural analyses in the present study did not suggest any handedness switching of the fibril filament. The helicity switching of the fibril filaments may require a force generated by MreB proteins applied in a specialized manner (*11, 27*).

The alignment of the MreB and fibril proteins in the internal ribbon is critical to understanding the roles of individual proteins in the swimming mechanism (*7, 8*). We succeeded in tracing the ribbon and visualizing the filamentous structures by cryo-ET of the *Spiroplasma* and syn3B cells (Fig. 4). Comparing the lateral filament intervals in the ribbon-like sheets visualized in the cells of *Spiroplasma*, syn3B with MreB4–5, and syn3B with fibril, we assigned the MreBs and fibril filaments to the central and peripheral parts of the internal ribbon. This conclusion is consistent with a suggestion based on an early cryo-EM study of *Spiroplasma melliferum* (*8*). We also succeeded in docking the atomic model of the isolated fibril filament to the density map of the ribbon of *Spiroplasma* cells obtained by subtomogram averaging (Fig. 5). However, we failed to dock the atomic model of the MreB filament to the density map (*28, 29*), because a higher resolution was needed than that in the case of the fibril filament.

In a previous study, swimming cells reconstituted by helicity switching were expressed by MreB4–5 in the syn3B cell (*14-16*). Moreover, it is known that a few *Spiroplasma* species can swim even without the fibril gene (*30*). This suggests that the fibril protein is necessary for neither cell helicity formation nor force generation. Therefore, the question of the role of the fibril protein in *Spiroplasma* remains. The fibril filament isolated from *Spiroplasma* cells showed helical parameters similar to those of *Spiroplasma* cells and syn3B cells expressing MreB4–5. Moreover, the fibril filament formed in the syn3B cells also showed identical helical parameters (Fig. 4b). Considering that the fibril protein forms stable and yet flexible filaments (Fig. 3) just beneath the cell membrane (Fig. 5).

These features suggest that the fibril protein may have a role in stabilizing the shape of the helical cell lacking the peptidoglycan layer to achieve the robustness and efficiency required for repetitive helicity switching of the cell shape.

In this study, we also succeeded in characterizing two types of filaments formed by MreBs and the fibril protein in the syn3B cells (Fig. 4b). The synthetic bacterium syn3B has been developed and used to study a minimal system for life (*31*). We have demonstrated that the simple cell structure and small thickness of syn3B may also be advantageous for the analysis of the internal cell structures by cryo-ET (Fig. 4b) (*32, 33*)

## Materials and Methods

### Sample preparation

JCVI-syn3B (GenBank: CP069345.1) (*15, 16*), its derivatives (*14*), and *Spiroplasma eriocheiris* (TDA-040725-5T) (*9, 34*) were cultured to mid-log phase in SP4 media at 37°C, 37°C, and 30°C, respectively. The promoter of syn3B for MreB4–5 expression was changed from Tuf to the native ones (Fig. S6). The fibril filament fraction was prepared as described previously and adjusted to a concentration of 1 mg/ml in a buffer consisting of 20 mM Tris-Cl pH 8.0 and 150 mM NaCl.

### Cryo-EM and image processing

A 1.4 μl sample solution was applied onto a glow-discharged holey carbon grid (Quantifoil R1.2/1.3 Mo grid, Quantifoil Micro Tools, Germany), and the grid was plunge-frozen into liquid ethane by Vitrobot mark IV (Thermo Fisher Science, USA) with a blotting time of 3 s at 18 °C and 90% humidity. All the data collection was performed on a prototype of the CRYO ARM 200 (JEOL, Japan) equipped with a thermal field-emission electron gun operated at 200 kV, an Ω-type energy filter with a 20 eV slit width, and a K2 Summit direct electron detector camera (Gatan, USA). An automated data acquisition program, JADAS (JEOL, Japan), was used to collect cryo-EM image data, and pre-processing, motion correction, and Contrast Transfer Function (CTF) estimation were carried out in real-time by the Gwatch image processing pipeline software (*35*). Movie frames were recorded using the K2 Summit camera at a calibrated magnification of ×40,000 corresponding to a pixel size of 1.5 Å with a defocus range from −0.6 to −1.8 μm. The data were collected with a total exposure of 10 s fractionated into 60 frames, with a total dose of ∼50 electrons Å^−2^ in counting mode. A total of 720 movies were collected.

Motion correction was performed by MotionCor2 v1.4.7, and CTF parameters were estimated by Gctf v1.06. Fibril filament images were automatically selected by RELION 3.0 as helical objects and segmented into 200 × 200-pixel boxes with 90% overlap. The best particles were selected from the results after the 2D classification of these segmented images by RELION 3.0. Segmented images of 237,690 were 3D classified into four classes, 147,040 segmented images from the best two classes were merged, and 3D auto-refine was applied. Atomic model construction was performed in COOT28, followed by real-space refinement using PHENIX29. All 3D maps and model images used in this paper were created with UCSF ChimeraX-1.3.

### Cryo-ET and image processing

A 3.0 μl sample solution containing 10 nm colloidal gold was applied onto a glow-discharged holey carbon grid (Quantifoil R1.2/1.3 Cu grid, Quantifoil Micro Tools, Germany). The grid was plunge-frozen into liquid ethane by Vitrobot mark IV (Thermo Fisher Science, USA) with a blotting time of 3 s at 4 °C and 90% humidity. All the data collection was performed on a CRYO ARM 300 II (JEOL, Japan) equipped with a cold field-emission electron gun operated at 300 kV, an Ω-type energy filter with a 20 eV slit width, and a K3 Summit direct electron detector camera (Gatan, USA). Movie frames were recorded using the K3 Summit camera at a calibrated magnification of ×25,000 corresponding to a pixel size of 2.343 Å with a defocus range from −0.6 to −1.8 μm. The data were acquired at a tilt range from −60 to 60 degrees with a dose-symmetric tilt scheme at 3° increments using serialEM 4.0.

Motion correction was performed by RELION’s implement, and CTF parameters were estimated by ctffind4. Tilt series alignment and tomogram generation were performed in IMOD 4.11. Subtomogram coordinates were manually selected by the 3dmod tool in IMOD. After 2D classification of the segmented subtomogram by RELION 4.0, the best set of particle images was selected from the results. 21,973 segmented subtomograms were 3D classified into three classes, 5,182 segmented subtomograms from the best classes were selected, and 3D auto-refine was performed. All 3D maps and model images used in this paper were created with UCSF ChimeraX.

## Acknowledgments

We thank Yuhei O Tahara, Daichi Takahashi, Ikuko Fujiwara, Takuma Toyonaga, Isil Tulum, Tasuku Hamaguchi, and Keisuke Kawakami at Osaka Metropolitan University, Japan for helpful discussions.

## Funding

MEXT KAKENHI Grants-in-Aid for Scientific Research A JP17H01544 (MM) JST CREST JPMJCR19S5 (MM)

the Osaka City University (OCU) Strategic Research Grant 2017 for top priority research (MM)

JSPS KAKENHI JP25000013 (KN)

the Platform Project for Supporting Drug Discovery and Life Science Research (BINDS) from AMED JP19am0101117 support number 1282 (KN)

Research Support Project for Life Science and Drug Discovery (Basis for Supporting Innovative Drug Discovery and Life Science Research (BINDS)) from AMED under Grant Number JP22ama121003 (KN)

the Cyclic Innovation for Clinical Empowerment (CiCLE) from AMED Grant Number JP17pc0101020 (KN)

JEOL YOKOGUSHI Research Alliance Laboratories of Osaka University (KN)

## Author contributions

Each author’s contribution(s) to the paper should be listed (we suggest following the CRediT model with each CRediT role given its line. No punctuation in the initials.

Conceptualization: YS, MM

Methodology: YS, TK, TM, HK, AK, FM, KN

Investigation: YS

Visualization: YS, TK, TM, AK, FM, KN

Supervision: KN, MM

Writing—original draft: YS, MM

Writing—review & editing: YS, TK, TM, HK, AK, FM, KN, MM

## Data and materials availability

All data were deposited to EMDATA (D_1300016148, D_1300037997).

