## Supplementary material for "Cryo-electron microscopy of cytoskeletal ‘fibril’ involved in *Spiroplasma* swimming": Fig.S1 Fig.S2 Fig.S3 Fig.S4 Fig.S5 Fig.S6

**This PDF file includes:**

Figs. S1 to S6

**Other Supplementary Materials for this manuscript include the following:**

Movies S1 to S2

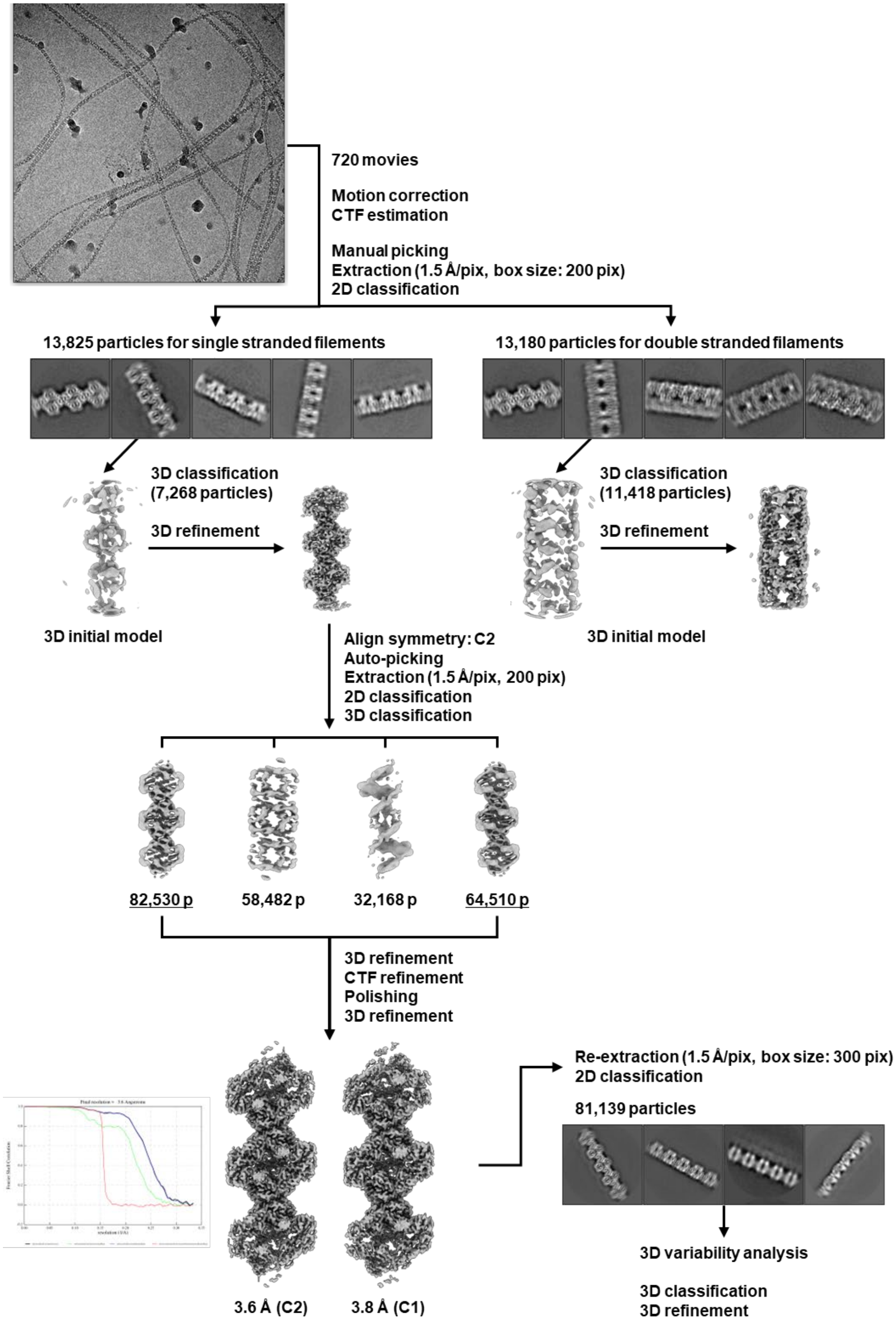


**Fig. S1. Single-particle analysis of fibril filament**

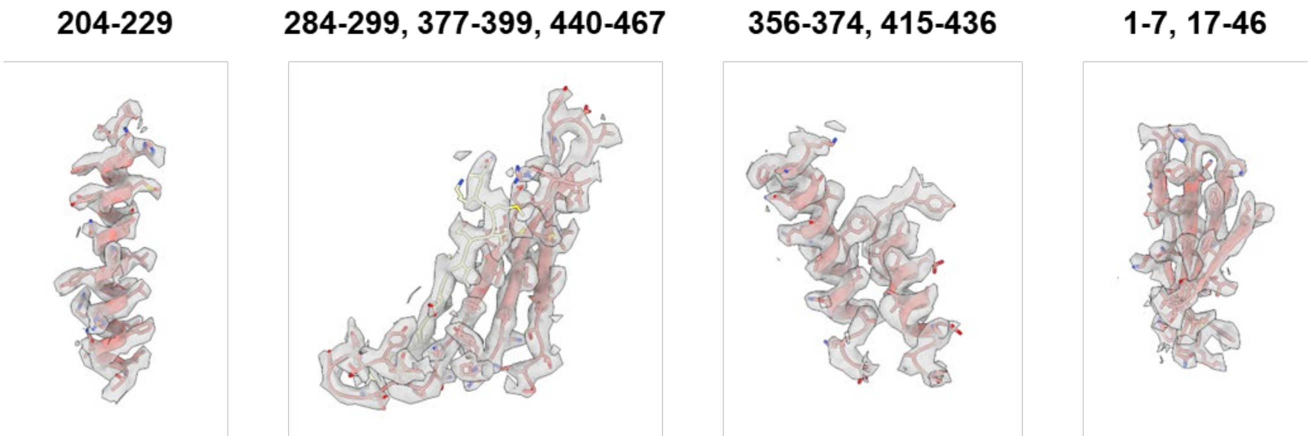


**Fig. S2. Cryo-EM maps and models of the fibril protein.**
The atomic model with representative maps at 3.6 Å resolution. The numbers indicate the amino acid residues.

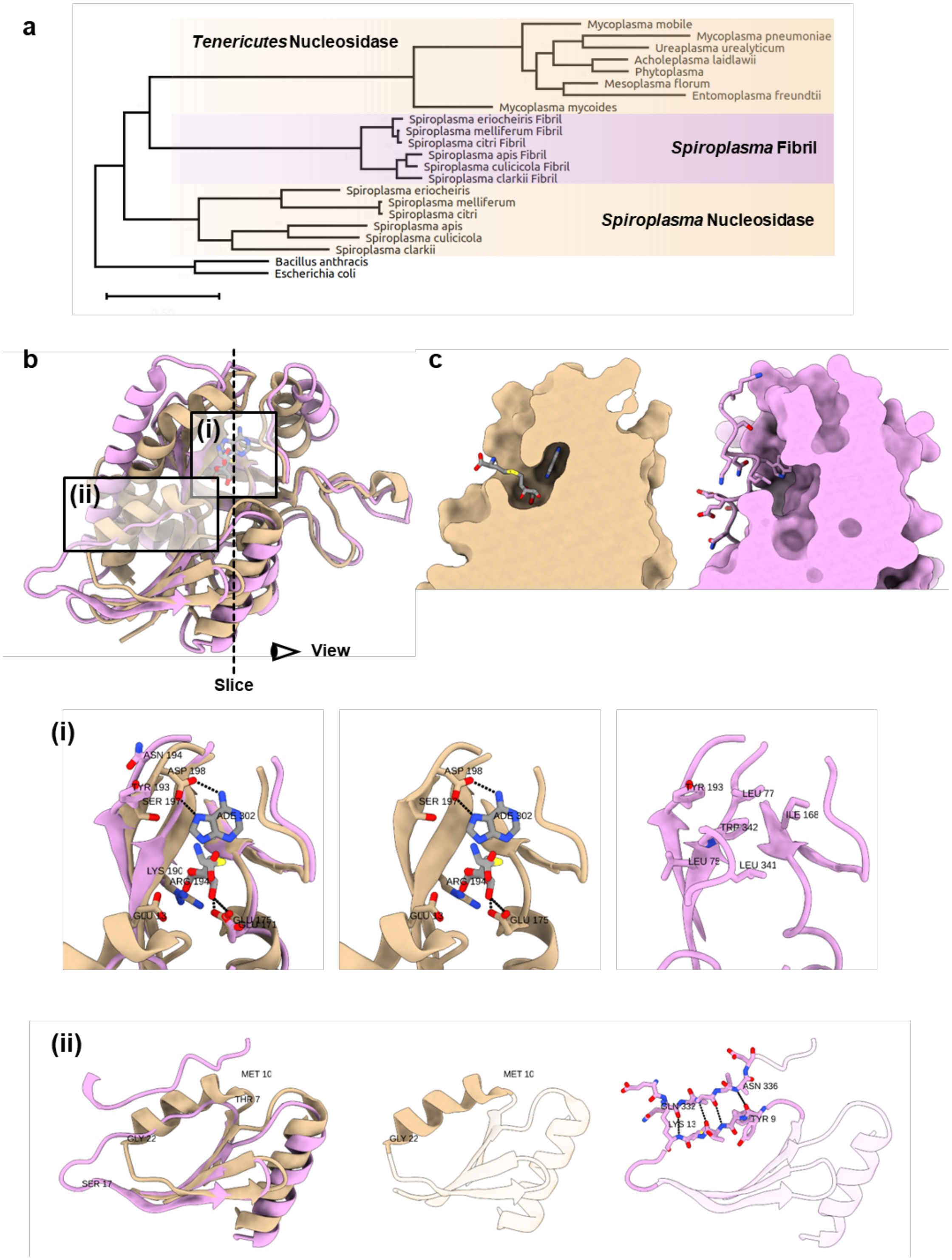


**Fig. 3S. Structural comparison between the fibril (pink) protein and bacterial SAH nucleosidase (beige). A** A phylogenetic tree including the nucleosidases and the fibril proteins. **B** Superposing of fibril protein and nucleosidase. The boxed areas (i) and (ii) are magnified below. **C** Comparison of binding pockets based on surface models.

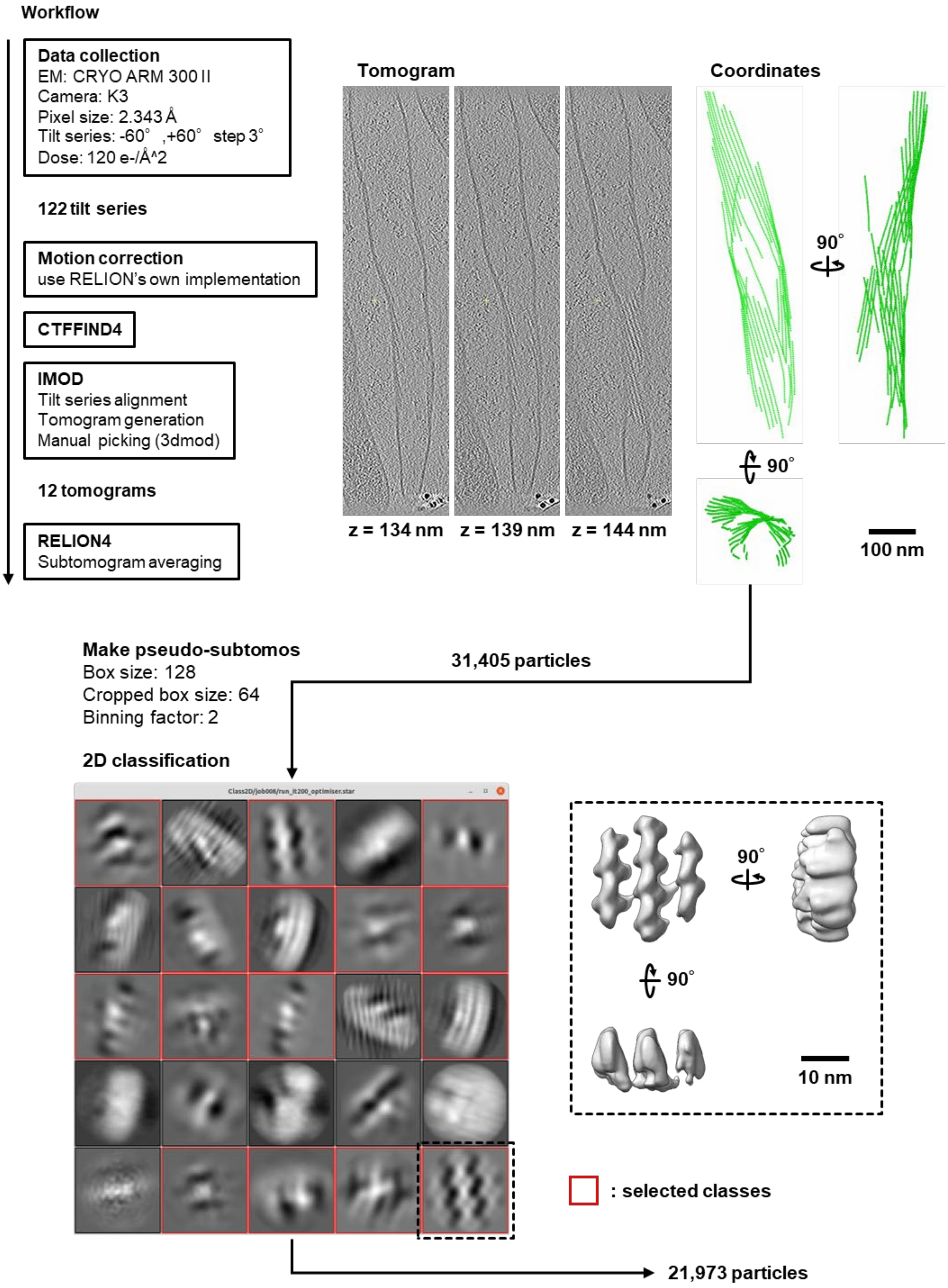


**Fig. S4-1. Workflow of cryo-ET and subtomogram averaging of the *in situ* fibril filament.**

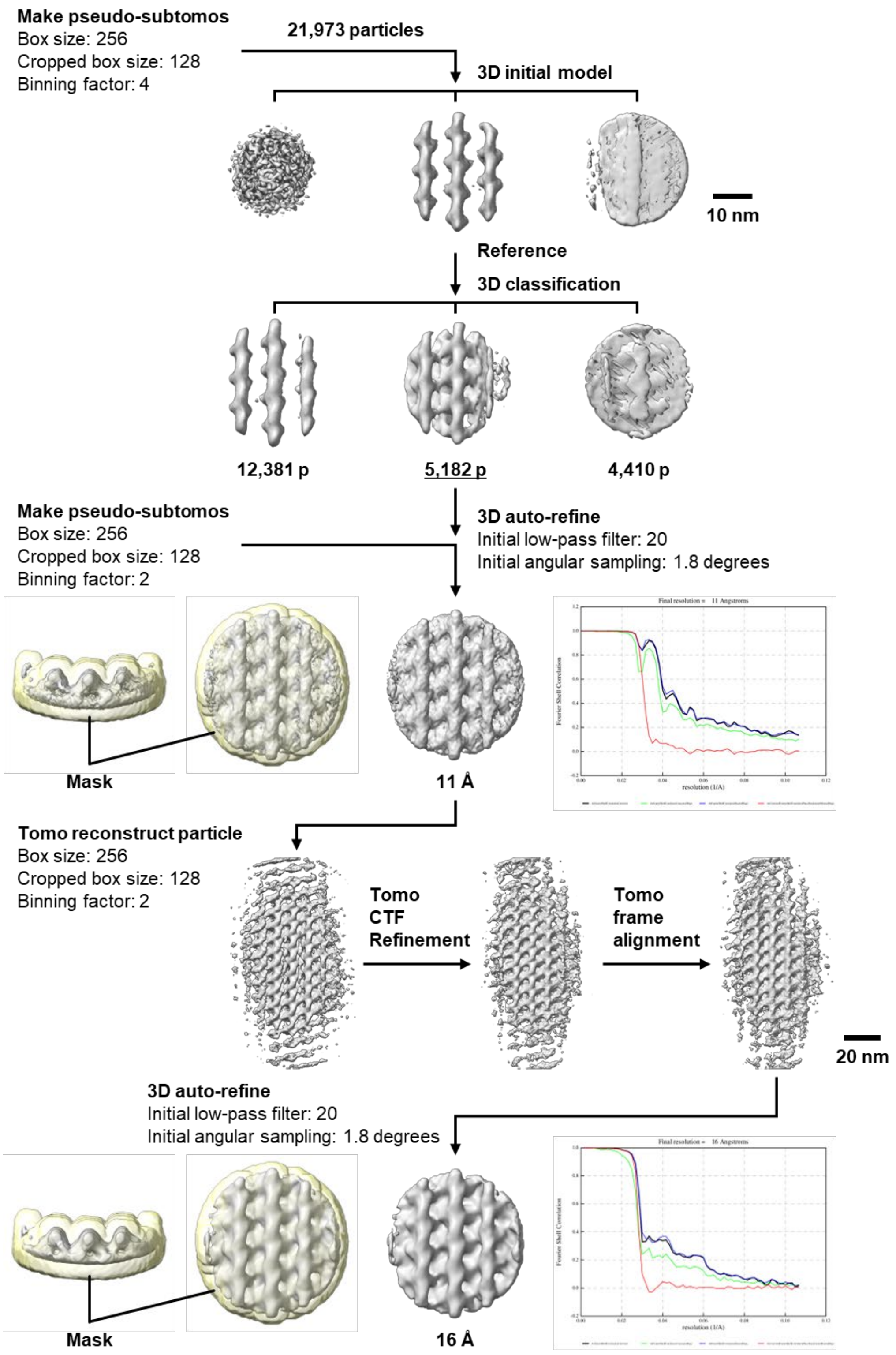


**Fig. S4-2. Workflow of cryo-ET and subtomogram averaging of the *in situ* fibril filament.**

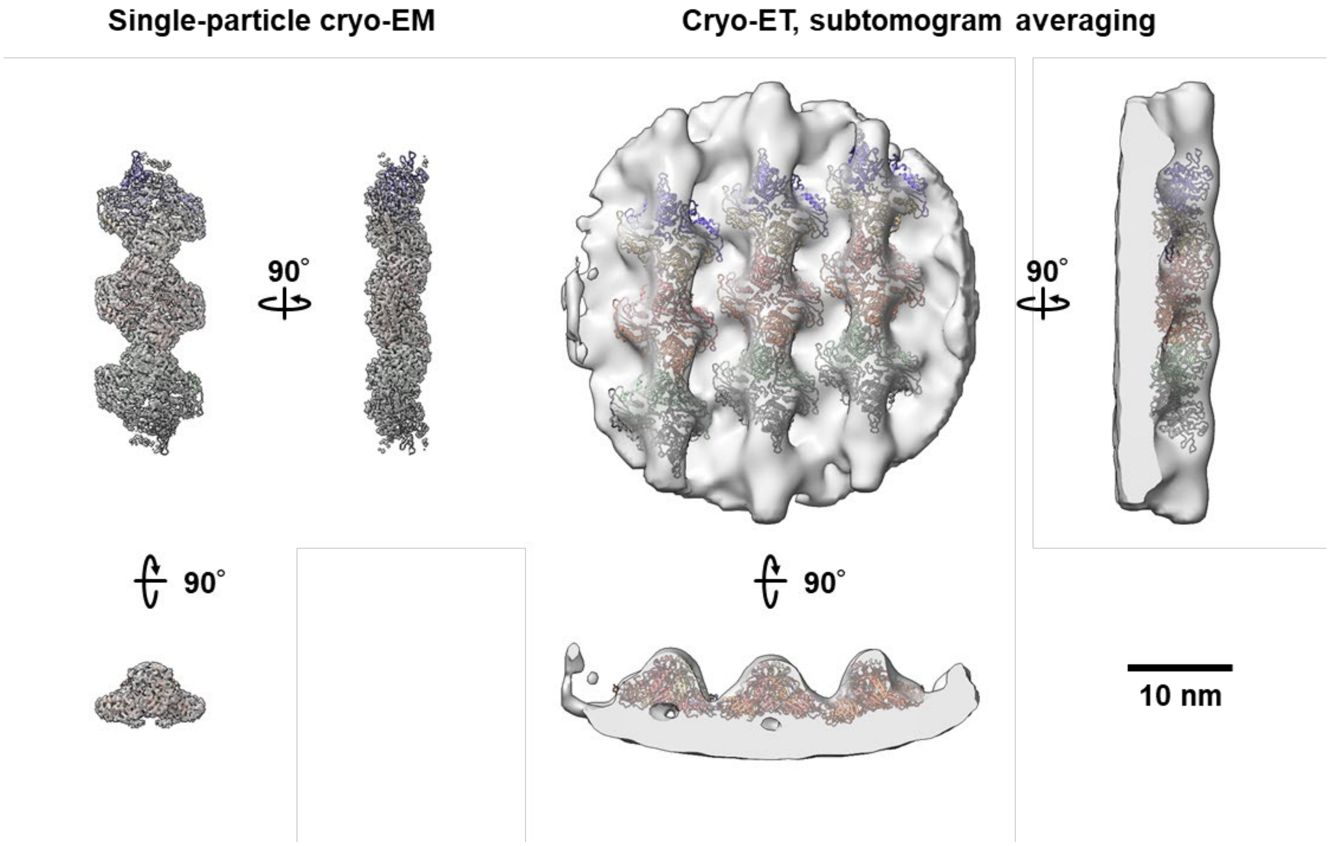


**Fig. S5. Fibril structures from single-particle analysis and subtomogram averaging.** The atomic structure was fitted to a structure from the subtomogram averaging. Each fibril subunit is colored differently.

**a** Ptuf–MreB4–MreB5

tattttttgaattaagtattaaataagtgtaaaatatataatagtaaaaacgccccaaaaggggcagacaaaatagtagaaatatatctattattcttgatttgttatagaaattttaa
ggagaaaaaaacatggcaggatttaatagcggcaaaagtaaaagaccaactttcgtttcaatggacttaggaactgcaaacacactagtgtatgtttcaggttcaggagta
gtttataacgaaccatcaatcgtagcttacagaataaaagaaaatagaattattgctgtagggatcgaagcttataaaatgatcggaaaaggtaacaaatcaattcgtattgta
agaccaatggttgacggagttattactgatattagagcaactgaagctcaattaagatatatcttcggaaaattacgaatctcaaaacaactaaaacactcaatcatgttattag
catgtccaagtgttattactgaattagaaaaagcagcgttgaaaaaaatcgcgatgaacttaggggcaactaaagttttcgttgaagaagaagttaaaatggctgccttaggt
ggaggagtagatatctacaaacctactggtaacttagttgttgatatgggagggggaacaacagatattgctgttattgcctcaggagatatcgtattatcgaaatcagtaaa
agttgctggaaactatctaaacgatgaaatgcaaaaattcattcgttcacaatatggattagaagtaggatcaaaaacagctgaacaaattaaaattgaaattggttcattagc
aaaatacccagacgaaagaaaaatgaaagtttatggacgtgacgtcgtttcaggattaccaagagaaatcgaagttacaccagaagaagttagagaggtattaaaagtac
cggtatcaagaattatcgacttaacagtccaagtattagaagaaacaccaccagaattggcaggagatatcttcaaaaatggaatcacaatttatggaggaggagcattaat
taagggaattgatcgttacttcacagatacattacaattaccatcaaaagttggtgaacaaccattactagcggttattaatggtactaaaaaattcgaatctgatatctatgaca
tcttacgtcaagaacaaatgcatactaaagaattagattactaaatattaaggaggaaattaacgtgaaaccagaaagaccatttatctcacttgacttgggaacagctaacg
tgttagcttacgtttctggtcaagggattatctataatgaaccatcattaatggcttacgatactaaaaccaataaattaattgctttagggaaagaagcttatgatatgattggaa
agacacacgaccaaattagaatggtgactccgctagttgatggagttatcgcagatatggaagctgcacaagatttattaaaacacgttttctcaagaatgaaaatgatgaat
atttgaaaaaatgcggtagttttactagcatgtccaagtggagttactgaattagaaagagaagctttaaaaaacgttgcccaagatatgggagctgacttagttattattgaa
gaagaagctaaaatggcagcaattggagcaggaatcaacattgacttaccacaagggaacttaatcattgatattggaggaggaacaactgacttagcaattatttcatcag
gagatattgtagttgcgagaagtattaaagttgccggaaaccactttgatgatgatatccgtaaatatattcgttcagaatataacattgcaattggacaaaaaactgcagaag
atgttaaaaaatacatcggttcattagttaaataccataatgaacgtgcaatgcaaatttatggaagagacattgtttcaggattaccaaaagaagctaaaattagttctgatga
aattagaaacgttttactaaatgccttttctaaaattactgacttagtaattgaattactggaaaatacacctccggaattagcaggagatatcatgcgtaatggtatcacaatttg
tggtgggggagcattaattagaaatattgataaatacttctttgatatcttccaattaccaacaagaactgcttctgatccgctaatgtgtgttattgaaggaacaagagcgtttg
aaaaagttattagaaaacgtattgaaaacggttactataacttcaatgacaaaggattattagctggcattgggaagaaaaaaataa

**b** Pnative–MreB4–MreB5

ctattgctaatttcaagaagtcattaattttttttattcaactataaaactctttatcaactaaaatgctagtaacatctcacttttaaaataatttttgttataataatttaaattaaattcttc
tagaagttctcttgcgaagaagattcctttttcttaaaatttatgttataattacttagtattgtatactctaatatcaatgtattagagtcttgcaatccatttttttactgacattaatagg
aaaattttttaatgttagatattgtttatgtttatacacattgaaaaagaaaaaagggaggaatttttacaatggcaggatttaatagcggcaaaagtaaaagaccaactttcgttt
caatggacttaggaactgcaaacacactagtgtatgtttcaggttcaggagtagtttataacgaaccatcaatcgtagcttacagaataaaagaaaatagaattattgctgtag
ggatcgaagcttataaaatgatcggaaaaggtaacaaatcaattcgtattgtaagaccaatggttgacggagttattactgatattagagcaactgaagctcaattaagatata
tcttcggaaaattacgaatctcaaaacaactaaaacactcaatcatgttattagcatgtccaagtgttattactgaattagaaaaagcagcgttgaaaaaaatcgcgatgaact
taggggcaactaaagttttcgttgaagaagaagttaaaatggctgccttaggtggaggagtagatatctacaaacctactggtaacttagttgttgatatgggagggggaac
aacagatattgctgttattgcctcaggagatatcgtattatcgaaatcagtaaaagttgctggaaactatctaaacgatgaaatgcaaaaattcattcgttcacaatatggatta
gaagtaggatcaaaaacagctgaacaaattaaaattgaaattggttcattagcaaaatacccagacgaaagaaaaatgaaagtttatggacgtgacgtcgtttcaggattac
caagagaaatcgaagttacaccagaagaagttagagaggtattaaaagtaccggtatcaagaattatcgacttaacagtccaagtattagaagaaacaccaccagaattg
gcaggagatatcttcaaaaatggaatcacaatttatggaggaggagcattaattaagggaattgatcgttacttcacagatacattacaattaccatcaaaagttggtgaaca
accattactagcggttattaatggtactaaaaaattcgaatctgatatctatgacatcttacgtcaagaacaaatgcatactaaagaattagattactaaaaaagaatatatttag
agaaattcgctttaattaaaaatatagaggtttttcttgatataatatttagagaatatattaaggaggaaattaacgtgaaaccagaaagaccatttatctcacttgacttggga
acagctaacgtgttagcttacgtttctggtcaagggattatctataatgaaccatcattaatggcttacgatactaaaaccaataaattaattgctttagggaaagaagcttatga
tatgattggaaagacacacgaccaaattagaatggtgactccgctagttgatggagttatcgcagatatggaagctgcacaagatttattaaaacacgttttctcaagaatga
aaatgatgaatatttgaaaaaatgcggtagttttactagcatgtccaagtggagttactgaattagaaagagaagctttaaaaaacgttgcccaagatatgggagctgactta
gttattattgaagaagaagctaaaatggcagcaattggagcaggaatcaacattgacttaccacaagggaacttaatcattgatattggaggaggaacaactgacttagca
attatttcatcaggagatattgtagttgcgagaagtattaaagttgccggaaaccactttgatgatgatatccgtaaatatattcgttcagaatataacattgcaattggacaaaa
aactgcagaagatgttaaaaaatacatcggttcattagttaaataccataatgaacgtgcaatgcaaatttatggaagagacattgtttcaggattaccaaaagaagctaaaat
tagttctgatgaaattagaaacgttttactaaatgccttttctaaaattactgacttagtaattgaattactggaaaatacacctccggaattagcaggagatatcatgcgtaatg
gtatcacaatttgtggtgggggagcattaattagaaatattgataaatacttctttgatatcttccaattaccaacaagaactgcttctgatccgctaatgtgtgttattgaaggaa
caagagcgtttgaaaaagttattagaaaacgtattgaaaacggttactataacttcaatgacaaaggattattagctggcattgggaagaaaaaaataattaacaagataaatta
aaaaaaacattaggttaacctaatgttttttttaatattaattcttaatccgttat

Fig. S6. Nucleotide sequences of mreB4–5 regions in syn3B constructs. **a** Original construct. **b** New construct. Modified regions are colored red.

**Movie S1.**
*Spiroplasma* cells reconstructed from cryo-ET.

**Movie S2.**
*Spiroplasma* and syn3 cells reconstructed from cryo-ET.
